# Crop-associated differences in soil chemical properties and root-associated bacterial communities between Welsh onion and sweet potato

**DOI:** 10.64898/2026.07.11.737990

**Authors:** Aiko Tanaka, Syunya Kubota, Takeshi Nakajima, Daigo Takemoto

**Affiliations:** Graduate School of Bioagricultural Sciences, Nagoya University, Furo-cho, Chikusa-ku, Nagoya, Aichi 464-8601, Japan; AGRI-CATION Co., Ltd., 2229-1 Kitayamada-cho, Kusatsu, Shiga 525-0061, Japan

**Keywords:** Welsh onion, sweet potato, root-associated bacteria, soil chemical properties, crop-associated microbiota

## Abstract

Crop species may shape soil chemical properties and root-associated microbiota, but direct comparisons between contrasting crops remain limited. We compared soils and root-associated bacterial communities of Welsh onion (*Allium fistulosum*) and sweet potato (*Ipomoea batatas*) under the same field context. Sweet potato soil showed significantly lower electrical conductivity, inorganic nitrogen, and Mg saturation than control soil. Root-associated communities differed between crops, whereas alpha diversity did not. Proteobacteria-related taxa were more represented in Welsh onion roots, whereas Actinomycetia-related taxa were more represented in sweet potato roots, providing a basis for future studies on crop-specific soil microbial management.

## Main text

Plants interact closely with diverse microbial communities in the rhizosphere, rhizoplane, and internal tissues. Root-associated bacterial communities are assembled from pre-existing soil microbial pools and are shaped by soil chemical properties, plant genotype, root traits, developmental stage, and environmental conditions; in turn, they may influence plant growth, nutrient acquisition, and stress tolerance (Trivedi et al., 2020). Roots recruit a subset of soil-derived bacterial communities, with Proteobacteria, Bacteroidetes/Bacteroidota, and Actinobacteria frequently detected as major root-associated groups in *Arabidopsis*, rice, and diverse plant species (Bulgarelli et al., 2012; Edwards et al., 2015; Fitzpatrick et al., 2018). Root exudates also provide important chemical cues and substrates that shape rhizosphere microbial communities (Huang et al., 2014). Therefore, comparisons between contrasting crops may provide insights into how crop-specific root traits are associated with soil chemical properties and root-associated bacterial communities.

Welsh onion (*Allium fistulosum*) and sweet potato (*Ipomoea batatas*) represent contrasting crop types. Welsh onion forms fibrous roots and is cultivated mainly for its aboveground tissues, whereas sweet potato develops storage roots and shows distinct belowground carbon allocation. To address how these contrasting crops are associated with root–soil environments, we compared soil chemical properties among uncultivated control soil, Welsh onion-cultivated soil, and sweet potato-cultivated soil within the same field in Kusatsu, Shiga, Japan, and further compared root-associated bacterial communities between Welsh onion and sweet potato.

We first compared soil chemical properties among uncultivated control, Welsh onion-cultivated, and sweet potato-cultivated soils. Soil samples were collected from control, Welsh onion, and sweet potato plots, with three independent samples per treatment. Soil pH and electrical conductivity (EC) were measured in a 1:5 soil:deionized water suspension using standard soil testing procedures (Soil Survey Staff, 2014). Total carbon, total nitrogen, exchangeable cations, cation exchange capacity, humus, and micronutrients were measured using standard soil chemical methods (Sparks et al., 1996). Ammonium nitrogen was determined by the indophenol method, nitrate nitrogen by alkaline reduction followed by diazotization colorimetry, and available phosphate by the Murphy–Riley molybdenum blue method (Murphy and Riley, 1962). Inorganic nitrogen was calculated as the sum of ammonium nitrogen and nitrate nitrogen. Unless otherwise stated, chemical properties were expressed on a dry soil basis. Soil chemical properties were compared using Welch’s t-test followed by Holm correction within each soil property.

Basic soil properties and organic matter-related parameters, including pH, bulk density, humus, total carbon, and total nitrogen, showed broadly overlapping ranges among the three soil types (Fig. 1A). In contrast, electrical conductivity (EC) was significantly lower in sweet potato soil than in control and Welsh onion-cultivated soil. Among nitrogen- and phosphorus-related parameters, inorganic nitrogen was also significantly lower in sweet potato soil (Fig. 1B). Ammonium nitrogen showed lower mean values in both Welsh onion and sweet potato soils, whereas nitrate nitrogen was particularly low in sweet potato soil. Available phosphate and phosphate absorption coefficient did not show clear decreases in Welsh onion and sweet potato soil.

**Fig. 1.**
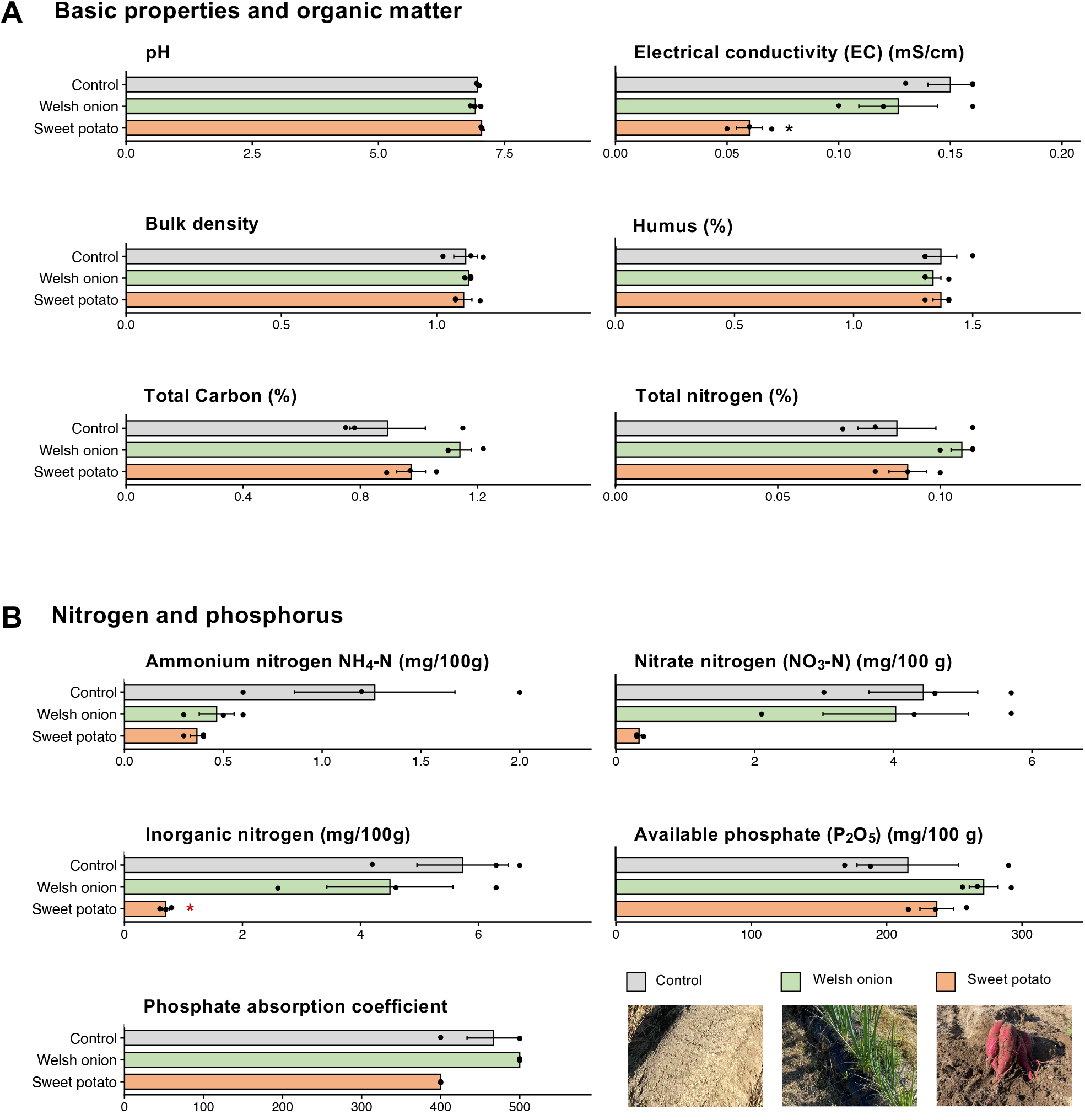
Soil chemical properties of control, Welsh onion, and sweet potato soils. **(A)** Basic soil properties and organic matter-related parameters, including pH, electrical conductivity (EC), bulk density, humus, total carbon, and total nitrogen. **(B)** Nitrogen- and phosphorus-related soil properties, including ammonium nitrogen, nitrate nitrogen, inorganic nitrogen, available phosphate, and phosphate absorption coefficient. Bars and error bars indicate means ± SE (n = 3), and black dots indicate individual samples. Asterisks indicate significant differences from the control by Welch’s t-test followed by Holm correction within each soil property (*P* < 0.05). Representative images of control, Welsh onion, and sweet potato soils are shown in the lower right of panel B.

Exchangeable cations, base balance, and micronutrient properties are summarized in Table 1. Sweet potato soil showed significantly lower Mg saturation than control soil. In addition, exchangeable K□O, exchangeable MgO, and K saturation showed lower mean values in sweet potato soil, suggesting a modest shift in exchangeable base-related properties. Among micronutrients, exchangeable Mn and soluble Zn also showed lower mean values in sweet potato soil, whereas water-soluble B, available Fe, and soluble Cu were not lower. Overall, these patterns suggest that sweet potato-cultivated soil was associated with lower values for selected nitrogen- and base-related properties, whereas Welsh onion-cultivated soil did not show a comparable downward shift relative to control soil.

**Table 1.** Exchangeable cations, base balance, and micronutrient properties of control, Welsh onion, and sweet potato soils.

| Soil property | Unit | Control | Welsh onion | Sweet potato |
| --- | --- | --- | --- | --- |
| Exchangeable K<sub>2</sub>O | mg/100 g | 18.0 ± 2.6 | 15.3 ± 0.9 | 10.0 ± 0.6 |
| Exchangeable CaO | mg/100 g | 203.0 ± 24.5 | 188.0 ± 9.5 | 193.3 ± 17.1 |
| Exchangeable MgO | mg/100 g | 20.7 ± 2.2 | 18.7 ± 0.3 | 12.0 ± 1.0 |
| Cation exchange capacity | meq/100 g | 7.8 ± 0.4 | 8.2 ± 0.2 | 7.2 ± 0.2 |
| K saturation | % | 4.9 ± 0.6 | 4.0 ± 0.1 | 2.9 ± 0.1 |
| Ca saturation | % | 92.3 ± 6.8 | 81.7 ± 5.2 | 96.3 ± 7.0 |
| Mg saturation | % | 13.3 ± 0.9 | 11.3 ± 0.3 | 8.0 ± 0.6* |
| Base saturation | % | 110.3 ± 7.8 | 96.7 ± 5.8 | 107.7 ± 6.6 |
| Ca/Mg equivalent ratio | equivalent ratio | 7.00 ± 0.12 | 7.37 ± 0.29 | 11.73 ± 1.63 |
| Mg/K equivalent ratio | equivalent ratio | 2.73 ± 0.30 | 2.80 ± 0.15 | 2.83 ± 0.12 |
| Water-soluble B | mg/kg | 0.50 ± 0.00 | 0.57 ± 0.03 | 0.63 ± 0.03 |
| Available Fe | mg/kg | 1.9 ± 0.0 | 2.2 ± 0.1 | 2.1 ± 0.1 |
| Exchangeable Mn | mg/kg | 0.7 ± 0.1 | 0.8 ± 0.2 | 0.5 ± 0.1 |
| Soluble Zn | mg/kg | 45.4 ± 6.2 | 56.9 ± 1.8 | 37.7 ± 0.6 |
| Soluble Cu | mg/kg | 1.43 ± 0.22 | 1.10 ± 0.15 | 1.87 ± 0.18 |

Values indicate means ± SE (n = 3). Asterisks indicate significant differences from the control by Welch's t-test followed by Holm correction within each soil property ( $P < 0.05$ ).

Root-associated bacterial communities were analyzed by shotgun metagenomics of gently washed Welsh onion and sweet potato roots, targeting tightly attached rhizoplane and putative endophytic fractions. Microbial cells were enriched from washed roots, and DNA was extracted using the DNeasy PowerSoil Pro Kit as described previously (Tanaka et al., 2025). Shotgun metagenomic libraries were paired-end sequenced. FASTQ reads were assembled using metaSPAdes, generating 1,364,749 contigs (Nurk et al., 2017). Assembled contigs were grouped into metagenomic bins using MetaBAT2 v1.7 for bin-level abundance analysis (Kang et al., 2019). Taxonomic classification of the bins was assigned using GTDB-Tk v2.3.2 (Chaumeil et al., 2022), and their relative abundance profiles across samples were calculated using CoverM (Aroney et al., 2025). Classified bins were further summarized at higher taxonomic levels for visualization and comparison. Bray–Curtis dissimilarities were calculated from bin-level relative abundance profiles and visualized by PCoA. Differentially represented bins were identified using Welch’s t-test on log_2_-transformed relative abundance values with a pseudocount, followed by Benjamini–Hochberg false discovery rate correction.

Bray–Curtis PCoA showed that Welsh onion and sweet potato samples were largely separated along PCoA1, which explained 82.7% of the variation (Fig. 2A). The effect size associated with crop identity was high (R^2^ = 0.818), although the exact permutation test did not reach conventional statistical significance (exact P = 0.100). Alpha diversity indices, including Shannon diversity, Simpson diversity, and Pielou evenness, showed similar ranges between the two crop species (Fig. 2B), suggesting that the observed crop-associated differences mainly reflected community composition rather than within-sample diversity.

**Fig. 2.**
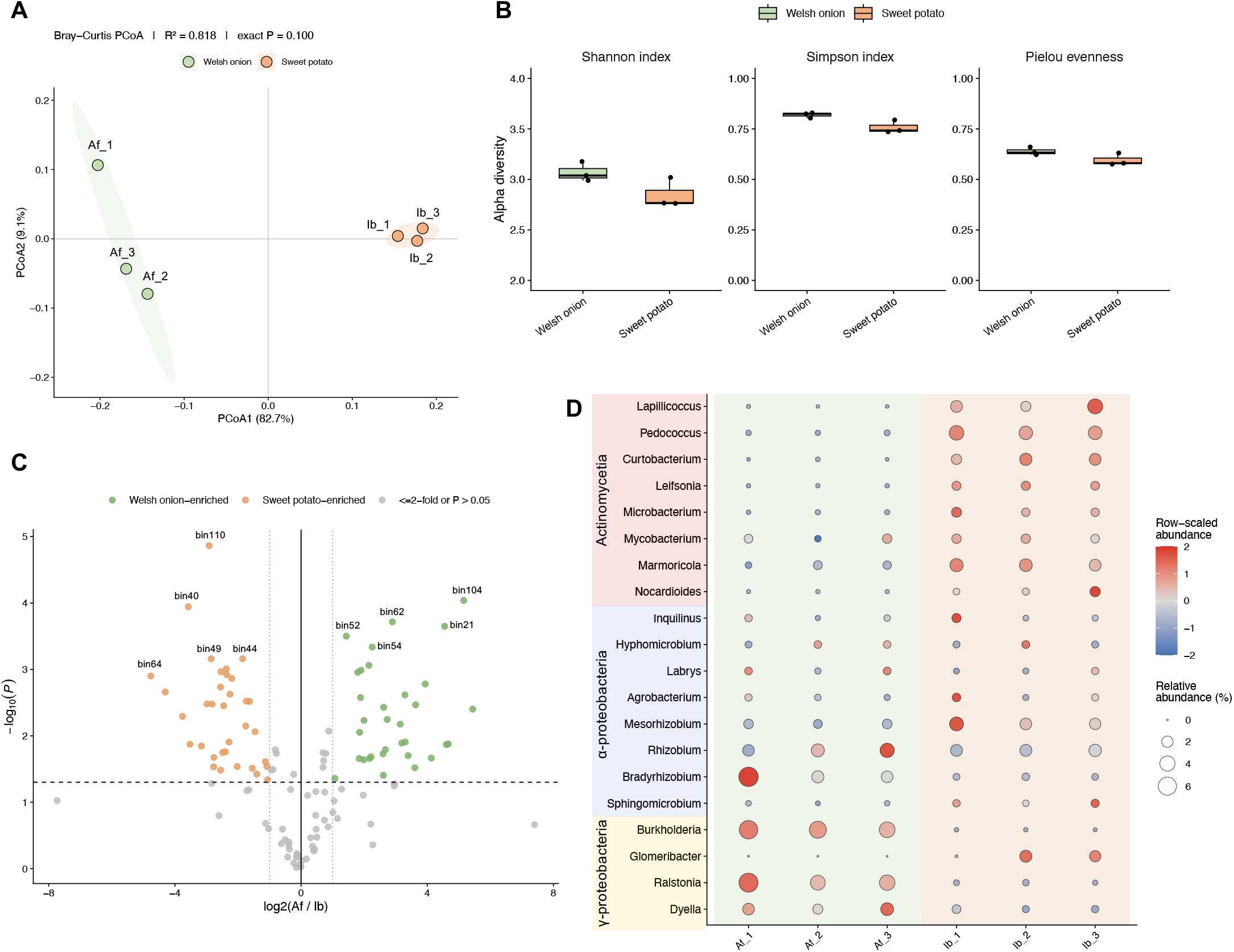
Comparison of root-associated bacterial communities between Welsh onion and sweet potato. **(A)** Principal coordinate analysis (PCoA) of root-associated bacterial communities. PCoA was performed using Bray–Curtis dissimilarities calculated from bin-level relative abundance profiles. Each point represents an independent root sample from Welsh onion (*Allium fistulosum*; Af_1–Af_3) or sweet potato (*Ipomoea batatas*; Ib_1–Ib_3). Shaded areas indicate group dispersion for visualization. Welsh onion and sweet potato samples were separated along PCoA1, which explained 82.7% of the variation (R^2^ = 0.818, exact *P* = 0.100). **(B)** Alpha diversity of root-associated bacterial communities. Shannon diversity, Simpson diversity, and Pielou evenness were calculated from bin-level relative abundance profiles. Box plots show the distribution of three independent samples per crop species, and black dots indicate individual samples. No statistically significant differences were detected between Welsh onion and sweet potato samples for these indices. **(C)** Volcano plot showing differentially represented bacterial bins between Welsh onion and sweet potato root samples. The x-axis indicates log_2_(Af/Ib), and the y-axis indicates -log_10_(P). P values were calculated using Welch’s t-test on log_2_-transformed relative abundance values with a pseudocount. Green and orange points indicate bins showing >2-fold higher relative abundance in Welsh onion and sweet potato samples, respectively, with *P* ≤ 0.05. Dotted lines indicate the thresholds of 2-fold difference and *P* = 0.05. **(D)** Bubble plot showing the relative abundance patterns of dominant classified bacterial genera. Unclassified bins were excluded before genus-level aggregation. Bubble size indicates relative abundance (%), and bubble color indicates row-scaled abundance across the six samples for each genus.

Differential abundance analysis of metagenomic bins further illustrated crop-associated differences in root-associated bacterial community profiles. The volcano plot showed multiple metagenomic bins with higher relative abundance in either Welsh onion or sweet potato samples (Fig. 2C). To place these bin-level patterns in a taxonomic context, classified bins were summarized at the genus level. Welsh onion samples showed higher representation of Proteobacteria-related genera, whereas sweet potato samples showed higher representation of Actinomycetia-related genera (Fig. 2D).

To further visualize representative metagenomic bins contributing to these crop-associated patterns, we examined bins with higher mean relative abundance in either Welsh onion or sweet potato samples (Fig. 3). Welsh onion-associated bins included classified representatives such as *Burkholderia, Bradyrhizobium, Ralstonia*, and *Dyella*. Sweet potato-associated bins included classified representatives such as *Marmoricola, Lapillicoccus, Pedococcus, Curtobacterium, Leifsonia*, and *Microbacterium*. These representative bins broadly mirrored the genus-level pattern in Fig. 2D.

**Fig. 3.**
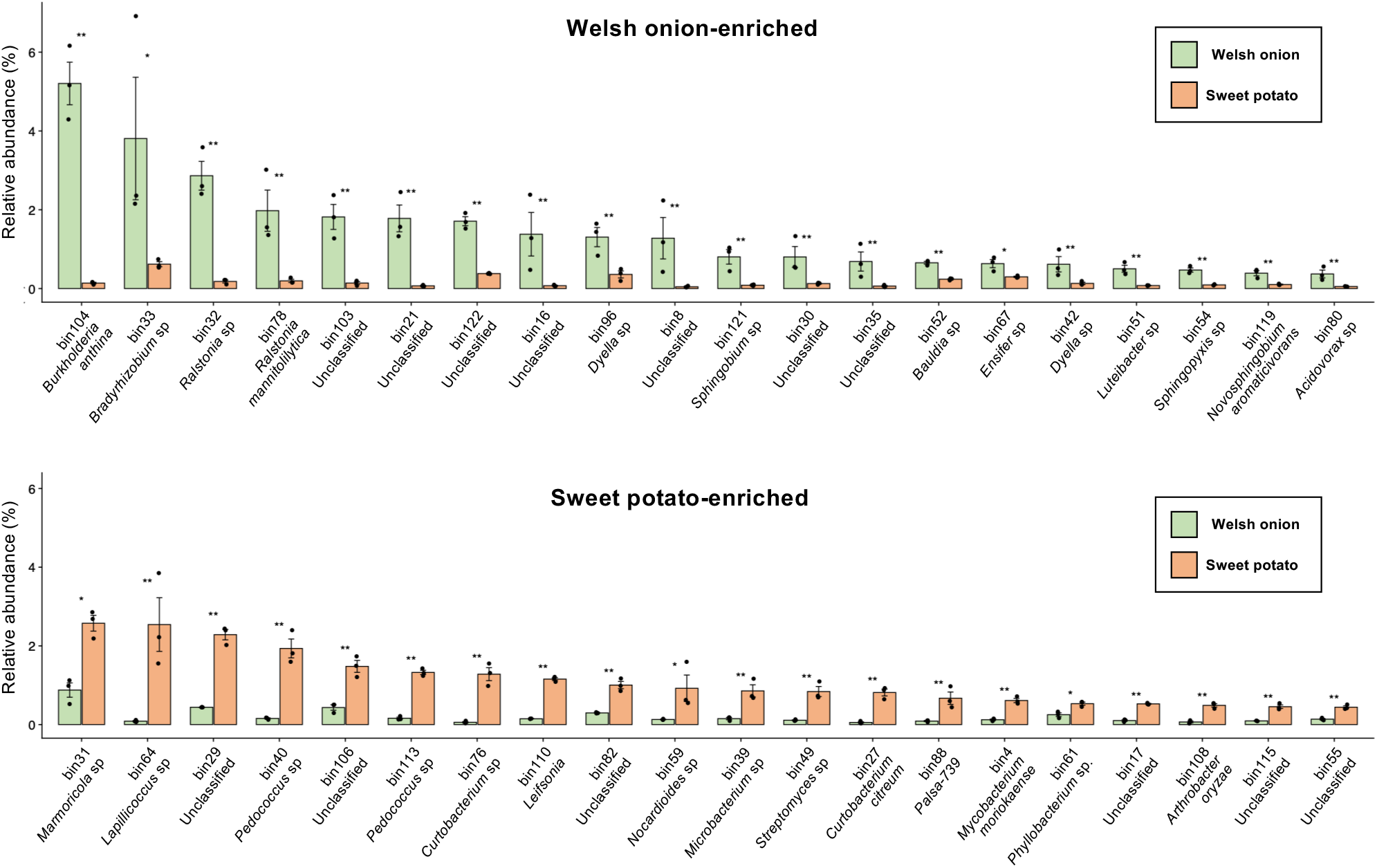
Representative bacterial bins showing crop-associated differences in root-associated bacterial communities. Relative abundances of representative bacterial bins that differed between Welsh onion and sweet potato root-associated bacterial communities are shown. The upper panel shows Welsh onion-enriched bins, ranked from left to right by mean relative abundance in Welsh onion samples. The lower panel shows sweet potato-enriched bins, ranked from left to right by mean relative abundance in sweet potato samples. Bins were selected based on >2-fold differences in mean relative abundance between crop species and nominal *P* < 0.05. Bars and error bars indicate means ± SE (n = 3), and black dots indicate individual samples. P values were calculated using Welch’s t-test on log_2_-transformed relative abundance values with a pseudocount, followed by Benjamini–Hochberg FDR correction. Double asterisks indicate FDR < 0.05, and single asterisks indicate nominal *P* < 0.05 but FDR ≥ 0.05.

Collectively, soil chemical and metagenomic analyses revealed crop-associated differences between Welsh onion and sweet potato under the same field context, with sweet potato-cultivated soil showing lower EC, inorganic nitrogen, Mg saturation, and several nutrient-related values. The contrasting representation of Proteobacteria-related and Actinomycetia-related groups is consistent with the general framework of root microbiome assembly, in which host identity, root traits, and rhizodeposition contribute to the recruitment of specific bacterial groups from soil-derived communities (Bulgarelli et al., 2012; Edwards et al., 2015; Huang et al., 2014; Fitzpatrick et al., 2018). Sweet potato is also notable because cultivated sweet potato carries *Agrobacterium*-derived cellular T-DNA, including *IbACS*, which encodes agrocinopine synthase (Kyndt et al., 2015; Tanaka et al., 2022). Tanaka et al. (2022) showed that *IbACS*-dependent agrocinopine A production enriched an actinobacterial *Leifsonia* strain in the tobacco root-associated/rhizosphere bacterial community. In the present study, sweet potato root-associated communities also showed higher representation of Actinomycetia-related groups, including *Leifsonia*. Although the present data do not establish a direct link between *IbACS* and these bacterial patterns, the results are consistent with the idea that sweet potato has crop-specific traits that may influence root-associated bacterial communities. By linking crop-associated soil chemistry with root-associated bacterial community structure, this study provides useful baseline information for understanding root–soil–microbe interactions and for guiding crop-specific soil microbial management.

## Acknowledgements

The authors thank Katakura & Co-op Agri Corporation, Tsukuba Analysis Center (Japan) for soil chemical analyses. Authors also thank Mr. Akira Ashida, Mr. Soichiro Egashira, Ms. Satomi Ohta and Ms. Akane Ueda (Nagoya University, Japan) and Mr. Wada (Otsu and Southern Shiga Agricultural Extension Division, Shiga Prefectural Government, Japan) for technical support. The authors used ChatGPT (OpenAI, GPT-5, 2025) only for English editing, and take full responsibility for the final text. This work was supported by Grants-in-Aid from the Japan Society for the Promotion of Science for Scientific Research (C) (JP22K06078 to A.T.) and Challenging Research (Exploratory) (JP22K19176 to D.T.).

## Supporting Information

**Supplementary Table S1**. Relative abundance and taxonomic classification of reconstructed metagenomic bins in Welsh onion and sweet potato root-associated bacterial communities.

## Notes

### Competing Interest Statement

The authors have declared no competing interest.

