## Supplemental Table S1 for "Crop-associated differences in soil chemical properties and root-associated bacterial communities between Welsh onion and sweet potato"

**Supplementary Table S1.** Relative abundance and taxonomic classification of reconstructed metagenomic bins in Welsh onion and sweet potato root-associated bacterial communities.

| | GTDB_classification1 | | | | | Welsh onion (*Allium fistulosum* ) | | | | | Sweet potato (*Ipomoea batatas* ) | | | | | | | | | | |
|---|---|---|---|---|---|---|---|---|---|---|---|---|---|---|---|---|---|---|---|---|---|
| Bin_ID | Class | Order | Family | Genus | Species | Af_1 | Af_2 | Af_3 | Mean (Af) | SE (Af) | Ib_1 | Ib_2 | Ib_3 | Mean (Ib) | SE (Ib) | Fold_change (Ib/Af)2 | Fold_change (Af/Ib)2 | Welch_P | BH_FDR | Higher_in | Significance |
| bin5 | Gammaproteobacteria | Burkholderiales | Burkholderiaceae | *Glomeribacter* | - | 0.000 | 0.000 | 0.000 | 0.000 | 0.000 | 0.023 | 2.339 | 1.999 | 1.453 | 0.722 | NA | 0.00 | 8.64E-02 | 1.21E-01 | | |
| bin64 | Actinomycetia | Actinomycetales | Dermatophilaceae | *Lapillicoccus* | - | 0.116 | 0.067 | 0.078 | 0.087 | 0.015 | 2.222 | 1.553 | 3.850 | 2.542 | 0.682 | 29.23 | 0.03 | 1.19E-03 | 9.22E-03 | Sweet potato | FDR < 0.05, P < 0.05 |
| bin76 | Actinomycetia | Actinomycetales | Microbacteriaceae | *Curtobacterium* | - | 0.039 | 0.096 | 0.040 | 0.058 | 0.019 | 0.982 | 1.552 | 1.310 | 1.281 | 0.165 | 22.00 | 0.05 | 2.42E-03 | 1.38E-02 | Sweet potato | FDR < 0.05, P < 0.05 |
| bin27 | Actinomycetia | Actinomycetales | Microbacteriaceae | *Curtobacterium* | *Curtobacterium citreum* | 0.033 | 0.091 | 0.038 | 0.054 | 0.019 | 0.644 | 0.877 | 0.926 | 0.815 | 0.087 | 15.11 | 0.07 | 5.53E-03 | 2.00E-02 | Sweet potato | FDR < 0.05, P < 0.05 |
| bin101 | Unclassified | - | - | - | - | 0.019 | 0.023 | 0.021 | 0.021 | 0.001 | 0.165 | 0.456 | 0.321 | 0.314 | 0.084 | 14.93 | 0.07 | 1.27E-02 | 3.50E-02 | Sweet potato | FDR < 0.05, P < 0.05 |
| bin40 | Actinomycetia | Actinomycetales | Dermatophilaceae | *Pedococcus* | - | 0.177 | 0.123 | 0.168 | 0.156 | 0.017 | 2.399 | 1.596 | 1.811 | 1.935 | 0.240 | 12.38 | 0.08 | 1.13E-04 | 4.63E-03 | Sweet potato | FDR < 0.05, P < 0.05 |
| bin11 | Actinomycetia | Actinomycetales | Micrococcaceae | *Arthrobacter* _E | *Arthrobacter woluwensis* | 0.025 | 0.036 | 0.029 | 0.030 | 0.003 | 0.456 | 0.161 | 0.344 | 0.320 | 0.086 | 10.76 | 0.09 | 1.34E-02 | 3.58E-02 | Sweet potato | FDR < 0.05, P < 0.05 |
| bin113 | Actinomycetia | Actinomycetales | Dermatophilaceae | *Pedococcus* | - | 0.130 | 0.139 | 0.216 | 0.162 | 0.027 | 1.239 | 1.427 | 1.311 | 1.326 | 0.055 | 8.21 | 0.12 | 3.38E-03 | 1.44E-02 | Sweet potato | FDR < 0.05, P < 0.05 |
| bin110 | Actinomycetia | Actinomycetales | Microbacteriaceae | *Leifsonia* | - | 0.146 | 0.144 | 0.152 | 0.148 | 0.002 | 1.170 | 1.211 | 1.087 | 1.156 | 0.036 | 7.84 | 0.13 | 1.29E-05 | 1.59E-03 | Sweet potato | FDR < 0.05, P < 0.05 |
| bin49 | Actinomycetia | Streptomycetales | Streptomycetaceae | *Streptomyces* | - | 0.132 | 0.093 | 0.107 | 0.111 | 0.012 | 1.096 | 0.692 | 0.732 | 0.840 | 0.128 | 7.60 | 0.13 | 6.64E-04 | 9.07E-03 | Sweet potato | FDR < 0.05, P < 0.05 |
| bin88 | Thermoleophilia | Gaiellales | Gaiellaceae | *Palsa* -739 | - | 0.060 | 0.107 | 0.101 | 0.089 | 0.015 | 0.599 | 0.973 | 0.440 | 0.671 | 0.158 | 7.52 | 0.13 | 3.22E-03 | 1.44E-02 | Sweet potato | FDR < 0.05, P < 0.05 |
| bin116 | Thermoleophilia | Solirubrobacterales | 70-9 | - | - | 0.064 | 0.127 | 0.213 | 0.135 | 0.043 | 1.820 | 0.295 | 0.903 | 1.006 | 0.443 | 7.46 | 0.13 | 5.15E-02 | 8.12E-02 | | |
| bin108 | Actinomycetia | Actinomycetales | Micrococcaceae | *Arthrobacter* _I | *Arthrobacter oryzae* | 0.029 | 0.108 | 0.062 | 0.066 | 0.023 | 0.403 | 0.517 | 0.544 | 0.488 | 0.043 | 7.39 | 0.14 | 2.23E-02 | 4.56E-02 | Sweet potato | FDR < 0.05, P < 0.05 |
| bin59 | Actinomycetia | Propionibacteriales | Nocardioidaceae | *Nocardioides* | - | 0.121 | 0.146 | 0.121 | 0.129 | 0.008 | 0.548 | 0.625 | 1.595 | 0.923 | 0.337 | 7.13 | 0.14 | 2.90E-02 | 5.56E-02 | Sweet potato | P < 0.05 |
| bin19 | Bacilli | Bacillales | Bacillaceae_H | *Priestia* | *Priestia megaterium* | 0.014 | 0.044 | 0.027 | 0.028 | 0.009 | 0.173 | 0.205 | 0.192 | 0.190 | 0.009 | 6.73 | 0.15 | 1.94E-02 | 4.41E-02 | Sweet potato | FDR < 0.05, P < 0.05 |
| bin75 | Chloroflexia | Thermomicrobiales | UBA6265 | - | - | 0.040 | 0.061 | 0.029 | 0.043 | 0.009 | 0.500 | 0.127 | 0.234 | 0.287 | 0.111 | 6.62 | 0.15 | 3.14E-02 | 5.60E-02 | Sweet potato | P < 0.05 |
| bin3 | Unclassified | - | - | - | - | 0.064 | 0.034 | 0.149 | 0.082 | 0.035 | 1.254 | 0.240 | 0.112 | 0.535 | 0.361 | 6.50 | 0.15 | 1.57E-01 | 2.10E-01 | | |
| bin66 | Actinomycetia | Mycobacteriales | Mycobacteriaceae | *Williamsia* _A | - | 0.043 | 0.082 | 0.052 | 0.059 | 0.012 | 0.261 | 0.426 | 0.452 | 0.380 | 0.060 | 6.42 | 0.16 | 1.88E-03 | 1.22E-02 | Sweet potato | FDR < 0.05, P < 0.05 |
| bin47 | Thermoleophilia | Gaiellales | Gaiellaceae | - | - | 0.045 | 0.080 | 0.069 | 0.065 | 0.010 | 0.354 | 0.530 | 0.345 | 0.410 | 0.060 | 6.33 | 0.16 | 1.13E-03 | 9.22E-03 | Sweet potato | FDR < 0.05, P < 0.05 |
| bin99 | Actinomycetia | Mycobacteriales | Mycobacteriaceae | *Smaragdicoccus* | - | 0.043 | 0.057 | 0.082 | 0.061 | 0.012 | 0.173 | 0.405 | 0.474 | 0.351 | 0.091 | 5.76 | 0.17 | 1.68E-02 | 4.21E-02 | Sweet potato | FDR < 0.05, P < 0.05 |
| bin39 | Actinomycetia | Actinomycetales | Microbacteriaceae | *Microbacterium* | - | 0.093 | 0.182 | 0.177 | 0.151 | 0.029 | 1.165 | 0.725 | 0.679 | 0.856 | 0.155 | 5.68 | 0.18 | 3.62E-03 | 1.44E-02 | Sweet potato | FDR < 0.05, P < 0.05 |
| bin85 | Actinomycetia | Streptosporangiales | Streptosporangiaceae | *Nonomuraea* | | 0.097 | 0.068 | 0.060 | 0.075 | 0.011 | 0.531 | 0.336 | 0.377 | 0.415 | 0.059 | 5.55 | 0.18 | 9.85E-04 | 9.22E-03 | Sweet potato | FDR < 0.05, P < 0.05 |
| bin29 | Unclassified | - | - | - | - | 0.438 | 0.438 | 0.446 | 0.440 | 0.003 | 2.395 | 2.438 | 2.023 | 2.285 | 0.132 | 5.19 | 0.19 | 1.20E-03 | 9.22E-03 | Sweet potato | FDR < 0.05, P < 0.05 |
| bin17 | Unclassified | - | - | - | - | 0.066 | 0.129 | 0.113 | 0.103 | 0.019 | 0.510 | 0.551 | 0.519 | 0.527 | 0.012 | 5.13 | 0.20 | 1.27E-02 | 3.50E-02 | Sweet potato | FDR < 0.05, P < 0.05 |
| bin4 | Actinomycetia | Mycobacteriales | Mycobacteriaceae | *Mycobacterium* | *Mycobacterium moriokaense* | 0.151 | 0.088 | 0.130 | 0.123 | 0.019 | 0.717 | 0.534 | 0.586 | 0.612 | 0.054 | 4.98 | 0.20 | 2.47E-03 | 1.38E-02 | Sweet potato | FDR < 0.05, P < 0.05 |
| bin115 | Actinomycetia | Mycobacteriales | Jatrophihabitantaceae | - | - | 0.086 | 0.098 | 0.097 | 0.094 | 0.004 | 0.430 | 0.543 | 0.390 | 0.455 | 0.046 | 4.85 | 0.21 | 1.31E-03 | 9.46E-03 | Sweet potato | FDR < 0.05, P < 0.05 |
| bin37 | Gammaproteobacteria | Burkholderiales | Burkholderiaceae | - | - | 0.067 | 0.098 | 0.027 | 0.064 | 0.021 | 0.301 | 0.361 | 0.181 | 0.281 | 0.053 | 4.39 | 0.23 | 3.06E-02 | 5.59E-02 | Sweet potato | P < 0.05 |
| bin44 | Actinomycetia | Actinomycetales | Microbacteriaceae | *Microbacterium* | - | 0.069 | 0.082 | 0.071 | 0.074 | 0.004 | 0.322 | 0.240 | 0.293 | 0.285 | 0.024 | 3.85 | 0.26 | 6.50E-04 | 9.07E-03 | Sweet potato | FDR < 0.05, P < 0.05 |
| bin26 | Alphaproteobacteria | Sphingomonadales | Sphingomonadaceae | *Sphingomicrobium* | - | 0.087 | 0.040 | 0.052 | 0.060 | 0.014 | 0.199 | 0.095 | 0.340 | 0.212 | 0.071 | 3.56 | 0.28 | 6.55E-02 | 9.71E-02 | | |
| bin106 | Unclassified | - | - | - | - | 0.294 | 0.506 | 0.500 | 0.434 | 0.070 | 1.734 | 1.494 | 1.208 | 1.479 | 0.152 | 3.41 | 0.29 | 7.19E-03 | 2.46E-02 | Sweet potato | FDR < 0.05, P < 0.05 |
| bin82 | Unclassified | - | - | - | - | 0.284 | 0.291 | 0.314 | 0.296 | 0.009 | 1.172 | 0.986 | 0.854 | 1.004 | 0.092 | 3.39 | 0.30 | 2.97E-03 | 1.44E-02 | Sweet potato | FDR < 0.05, P < 0.05 |
| bin22 | Alphaproteobacteria | Sphingomonadales | Sphingomonadaceae | *Sphingomicrobium* | - | 0.098 | 0.045 | 0.060 | 0.067 | 0.016 | 0.216 | 0.106 | 0.357 | 0.226 | 0.073 | 3.36 | 0.30 | 6.41E-02 | 9.66E-02 | | |
| bin55 | Thermoleophilia | Solirubrobacterales | 70-9 | - | - | 0.133 | 0.171 | 0.110 | 0.138 | 0.018 | 0.418 | 0.398 | 0.510 | 0.442 | 0.034 | 3.20 | 0.31 | 3.12E-03 | 1.44E-02 | Sweet potato | FDR < 0.05, P < 0.05 |
| bin14 | Actinomycetia | Mycobacteriales | Mycobacteriaceae | *Nocardia* | *Nocardia niigatensis* | 0.054 | 0.077 | 0.060 | 0.064 | 0.007 | 0.166 | 0.249 | 0.148 | 0.188 | 0.031 | 2.94 | 0.34 | 8.37E-03 | 2.71E-02 | Sweet potato | FDR < 0.05, P < 0.05 |
| bin31 | Actinomycetia | Propionibacteriales | Nocardioidaceae | *Marmoricola* | - | 0.522 | 1.125 | 0.985 | 0.877 | 0.182 | 2.856 | 2.685 | 2.188 | 2.576 | 0.201 | 2.94 | 0.34 | 3.09E-02 | 5.59E-02 | Sweet potato | P < 0.05 |
| bin79 | Unclassified | - | - | - | - | 0.053 | 0.042 | 0.041 | 0.045 | 0.004 | 0.092 | 0.109 | 0.193 | 0.132 | 0.031 | 2.90 | 0.35 | 3.65E-02 | 6.14E-02 | Sweet potato | P < 0.05 |
| bin63 | Unclassified | - | - | - | - | 0.058 | 0.106 | 0.090 | 0.085 | 0.014 | 0.222 | 0.181 | 0.178 | 0.194 | 0.014 | 2.29 | 0.44 | 2.52E-02 | 4.99E-02 | Sweet potato | FDR < 0.05, P < 0.05 |
| bin117 | Alphaproteobacteria | Rhizobiales_A | Rhizobiaceae_A | *Ochrobactrum* | *Ochrobactrum intermedium* | 0.242 | 0.096 | 0.243 | 0.193 | 0.049 | 0.766 | 0.219 | 0.304 | 0.430 | 0.170 | 2.22 | 0.45 | 2.07E-01 | 2.69E-01 | | |
| bin12 | Alphaproteobacteria | Rhizobiales | Devosiaceae | *Devosia* | - | 0.200 | 0.106 | 0.142 | 0.149 | 0.027 | 0.364 | 0.344 | 0.262 | 0.323 | 0.031 | 2.17 | 0.46 | 2.94E-02 | 5.56E-02 | Sweet potato | P < 0.05 |
| bin1 | Saccharimonadia | Saccharimonadales | UBA4665 | PMNU01 | - | 0.083 | 0.031 | 0.190 | 0.101 | 0.047 | 0.279 | 0.274 | 0.088 | 0.214 | 0.063 | 2.11 | 0.47 | 2.52E-01 | 3.10E-01 | | |
| bin61 | Alphaproteobacteria | Rhizobiales_A | Rhizobiaceae_A | *Phyllobacterium* | *Phyllobacterium* sp. 900539805 | 0.263 | 0.327 | 0.164 | 0.251 | 0.047 | 0.451 | 0.577 | 0.566 | 0.531 | 0.040 | 2.11 | 0.47 | 4.61E-02 | 7.46E-02 | Sweet potato | P < 0.05 |
| bin58 | Alphaproteobacteria | Rhizobiales | Rhizobiaceae | *Mesorhizobium* | - | 1.368 | 1.125 | 1.376 | 1.290 | 0.082 | 3.249 | 2.171 | 2.039 | 2.486 | 0.383 | 1.93 | 0.52 | 3.25E-02 | 5.63E-02 | Sweet potato | P < 0.05 |
| bin7 | Alphaproteobacteria | Rhizobiales | Methyloligellaceae | *Methyloceanibacter* | | 0.160 | 0.237 | 0.147 | 0.181 | 0.028 | 0.284 | 0.434 | 0.307 | 0.341 | 0.047 | 1.89 | 0.53 | 3.22E-02 | 5.63E-02 | Sweet potato | P < 0.05 |
| bin69 | Actinomycetia | Mycobacteriales | Mycobacteriaceae | *Mycobacterium* | *Mycobacterium* sp. 900240945 | 0.201 | 0.137 | 0.177 | 0.172 | 0.019 | 0.334 | 0.324 | 0.265 | 0.308 | 0.021 | 1.79 | 0.56 | 1.64E-02 | 4.20E-02 | Sweet potato | FDR < 0.05, P < 0.05 |
| bin34 | Alphaproteobacteria | Sphingomonadales | Sphingomonadaceae | *Sphingomicrobium* | - | 0.142 | 0.129 | 0.180 | 0.150 | 0.015 | 0.295 | 0.284 | 0.211 | 0.264 | 0.027 | 1.75 | 0.57 | 1.84E-02 | 4.27E-02 | Sweet potato | FDR < 0.05, P < 0.05 |
| bin68 | Babeliae | Babeliales | - | - | - | 0.036 | 0.091 | 0.101 | 0.076 | 0.020 | 0.053 | 0.125 | 0.183 | 0.120 | 0.038 | 1.59 | 0.63 | 4.22E-01 | 4.90E-01 | | |
| bin83 | Gammaproteobacteria | Xanthomonadales | Rhodanobacteraceae | *Dyella* | *Dyella jiangningensis* | 0.168 | 0.167 | 0.291 | 0.209 | 0.041 | 0.426 | 0.334 | 0.192 | 0.317 | 0.068 | 1.52 | 0.66 | 2.54E-01 | 3.10E-01 | | |
| bin112 | Unclassified | - | - | - | - | 0.036 | 0.130 | 0.149 | 0.105 | 0.035 | 0.114 | 0.175 | 0.167 | 0.152 | 0.019 | 1.45 | 0.69 | 3.68E-01 | 4.36E-01 | | |
| bin91 | Alphaproteobacteria | DSM-16000 | Inquilinaceae | *Inquilinus* | - | 0.725 | 0.269 | 0.489 | 0.494 | 0.132 | 1.215 | 0.362 | 0.422 | 0.666 | 0.275 | 1.35 | 0.74 | 6.68E-01 | 7.06E-01 | | |
| bin9 | Unclassified | - | - | - | - | 0.038 | 0.135 | 0.147 | 0.107 | 0.035 | 0.101 | 0.158 | 0.166 | 0.142 | 0.020 | 1.33 | 0.75 | 4.45E-01 | 5.11E-01 | | |
| bin45 | Actinomycetia | Propionibacteriales | Propionibacteriaceae | Fen-668 | - | 0.126 | 0.238 | 0.474 | 0.279 | 0.103 | 0.320 | 0.364 | 0.415 | 0.366 | 0.027 | 1.31 | 0.76 | 4.00E-01 | 4.69E-01 | | |
| bin105 | Alphaproteobacteria | Rhizobiales | Rhizobiaceae | *Agrobacterium* | *Agrobacterium pusense* | 0.694 | 0.498 | 0.420 | 0.537 | 0.081 | 1.021 | 0.419 | 0.654 | 0.698 | 0.175 | 1.30 | 0.77 | 5.10E-01 | 5.68E-01 | | |
| bin10 | Alphaproteobacteria | Rhizobiales | Hyphomicrobiaceae | *Hyphomicrobium* | - | 0.284 | 0.362 | 0.316 | 0.321 | 0.023 | 0.370 | 0.452 | 0.394 | 0.405 | 0.024 | 1.26 | 0.79 | 6.44E-02 | 9.66E-02 | | |
| bin72 | Babeliae | Babeliales | Vermiphilaceae | - | - | 0.075 | 0.094 | 0.075 | 0.081 | 0.006 | 0.046 | 0.104 | 0.141 | 0.097 | 0.028 | 1.19 | 0.84 | 8.26E-01 | 8.47E-01 | | |
| bin95 | Actinomycetia | Mycobacteriales | Mycobacteriaceae | *Nocardia* | *Nocardia elegans* | 0.257 | 0.338 | 0.269 | 0.288 | 0.025 | 0.514 | 0.264 | 0.180 | 0.319 | 0.100 | 1.11 | 0.90 | 9.60E-01 | 9.60E-01 | | |
| bin41 | Gammaproteobacteria | Enterobacterales | Enterobacteriaceae | *Enterobacter* | *Enterobacter asburiae* | 0.121 | 0.290 | 0.382 | 0.264 | 0.077 | 0.392 | 0.248 | 0.237 | 0.293 | 0.050 | 1.11 | 0.90 | 6.72E-01 | 7.06E-01 | | |
| bin118 | Alphaproteobacteria | Acetobacterales | Acetobacteraceae | *Roseomonas* _B | - | 0.409 | 0.240 | 0.244 | 0.298 | 0.056 | 0.351 | 0.269 | 0.363 | 0.328 | 0.029 | 1.10 | 0.91 | 5.92E-01 | 6.44E-01 | | |
| bin20 | Unclassified | - | - | - | - | 0.256 | 0.201 | 0.155 | 0.204 | 0.029 | 0.203 | 0.194 | 0.270 | 0.222 | 0.024 | 1.09 | 0.92 | 6.20E-01 | 6.69E-01 | | |
| bin114 | Unclassified | - | - | - | - | 0.603 | 0.609 | 0.657 | 0.623 | 0.017 | 0.799 | 0.587 | 0.598 | 0.661 | 0.069 | 1.06 | 0.94 | 6.70E-01 | 7.06E-01 | | |
| bin2 | Unclassified | - | - | - | - | 0.256 | 0.144 | 0.174 | 0.191 | 0.034 | 0.250 | 0.150 | 0.192 | 0.197 | 0.029 | 1.03 | 0.97 | 8.73E-01 | 8.88E-01 | | |
| bin73 | Unclassified | - | - | - | - | 0.128 | 0.227 | 0.236 | 0.197 | 0.035 | 0.243 | 0.243 | 0.106 | 0.197 | 0.046 | 1.00 | 1.00 | 9.38E-01 | 9.45E-01 | | |
| bin92 | Gammaproteobacteria | UBA6186 | UBA6186 | CADEGZ01 | - | 0.029 | 0.157 | 0.071 | 0.085 | 0.038 | 0.079 | 0.099 | 0.073 | 0.084 | 0.008 | 0.98 | 1.02 | 7.49E-01 | 7.74E-01 | | |
| bin98 | Alphaproteobacteria | Rhizobiales | Rhizobiaceae | *Rhizobium* | *Rhizobium* sp. 900469475 | 1.569 | 2.330 | 3.045 | 2.315 | 0.426 | 1.932 | 2.066 | 2.182 | 2.060 | 0.072 | 0.89 | 1.12 | 7.15E-01 | 7.45E-01 | | |
| bin24 | Unclassified | - | - | - | - | 0.327 | 0.284 | 0.279 | 0.297 | 0.015 | 0.272 | 0.252 | 0.243 | 0.256 | 0.008 | 0.86 | 1.16 | 7.89E-02 | 1.12E-01 | | |
| bin60 | Alphaproteobacteria | Rhizobiales | Hyphomicrobiaceae | *Hyphomicrobium* | - | 0.178 | 0.242 | 0.274 | 0.231 | 0.028 | 0.157 | 0.254 | 0.147 | 0.186 | 0.034 | 0.80 | 1.24 | 3.43E-01 | 4.09E-01 | | |
| bin6 | Alphaproteobacteria | Rhizobiales | Rhizobiaceae | *Allorhizobium* | - | 0.123 | 0.193 | 0.171 | 0.162 | 0.021 | 0.222 | 0.035 | 0.117 | 0.125 | 0.054 | 0.77 | 1.30 | 4.59E-01 | 5.22E-01 | | |
| bin13 | Saccharimonadia | Saccharimonadales | Saccharimonadaceae | | | 0.108 | 0.370 | 0.275 | 0.251 | 0.076 | 0.375 | 0.065 | 0.124 | 0.188 | 0.095 | 0.75 | 1.33 | 5.39E-01 | 5.92E-01 | | |
| bin53 | Saccharimonadia | Saccharimonadales | - | - | - | 0.132 | 0.401 | 0.301 | 0.278 | 0.078 | 0.380 | 0.086 | 0.149 | 0.205 | 0.089 | 0.74 | 1.36 | 5.12E-01 | 5.68E-01 | | |
| bin97 | Acidimicrobiia | Acidimicrobiales | Ilumatobacteraceae | - | - | 0.313 | 0.191 | 0.260 | 0.255 | 0.035 | 0.170 | 0.184 | 0.195 | 0.183 | 0.007 | 0.72 | 1.39 | 1.57E-01 | 2.10E-01 | | |
| bin18 | Unclassified | - | - | - | - | 0.469 | 0.341 | 0.374 | 0.395 | 0.039 | 0.339 | 0.253 | 0.258 | 0.283 | 0.028 | 0.72 | 1.39 | 6.90E-02 | 1.01E-01 | | |
| bin87 | Alphaproteobacteria | Rhizobiales | Labraceae | *Labrys* | - | 0.781 | 0.461 | 0.804 | 0.682 | 0.111 | 0.373 | 0.418 | 0.671 | 0.487 | 0.093 | 0.71 | 1.40 | 2.55E-01 | 3.10E-01 | | |
| bin46 | Unclassified | - | - | - | - | 0.721 | 1.305 | 1.580 | 1.202 | 0.253 | 0.841 | 0.858 | 0.848 | 0.849 | 0.005 | 0.71 | 1.42 | 3.37E-01 | 4.06E-01 | | |
| bin109 | Unclassified | - | - | - | - | 1.154 | 0.858 | 0.886 | 0.966 | 0.094 | 0.691 | 0.552 | 0.543 | 0.595 | 0.048 | 0.62 | 1.62 | 1.80E-02 | 4.27E-02 | Welsh onion | FDR < 0.05, P < 0.05 |
| bin70 | Alphaproteobacteria | Rhizobiales | Rhizobiaceae | *Rhizobium* | *Rhizobium* sp. 900471515 | 0.567 | 0.483 | 0.399 | 0.483 | 0.048 | 0.300 | 0.240 | 0.335 | 0.292 | 0.028 | 0.60 | 1.66 | 2.35E-02 | 4.75E-02 | Welsh onion | FDR < 0.05, P < 0.05 |
| bin111 | Alphaproteobacteria | Rhizobiales | Hyphomicrobiaceae | *Hyphomicrobium* | *Hyphomicrobium* sp. 003987855 | 0.153 | 0.169 | 0.155 | 0.159 | 0.005 | 0.088 | 0.116 | 0.082 | 0.096 | 0.010 | 0.60 | 1.66 | 3.01E-02 | 5.59E-02 | Welsh onion | P < 0.05 |
| bin56 | Unclassified | - | - | - | - | 0.185 | 0.350 | 0.409 | 0.315 | 0.067 | 0.184 | 0.195 | 0.177 | 0.185 | 0.005 | 0.59 | 1.70 | 1.86E-01 | 2.44E-01 | | |
| bin102 | Unclassified | - | - | - | - | 0.398 | 0.238 | 0.257 | 0.298 | 0.050 | 0.227 | 0.148 | 0.149 | 0.175 | 0.026 | 0.59 | 1.70 | 7.03E-02 | 1.02E-01 | | |
| bin25 | Unclassified | - | - | - | - | 0.228 | 0.166 | 0.233 | 0.209 | 0.022 | 0.129 | 0.123 | 0.103 | 0.118 | 0.008 | 0.57 | 1.77 | 1.83E-02 | 4.27E-02 | Welsh onion | FDR < 0.05, P < 0.05 |
| bin77 | Unclassified | - | - | - | - | 0.390 | 1.080 | 0.863 | 0.777 | 0.203 | 0.446 | 0.434 | 0.399 | 0.426 | 0.014 | 0.55 | 1.82 | 2.35E-01 | 2.95E-01 | | |
| bin94 | Alphaproteobacteria | Rhizobiales | Rhizobiaceae | FKL33 | - | 0.238 | 0.331 | 0.271 | 0.280 | 0.027 | 0.171 | 0.138 | 0.140 | 0.150 | 0.011 | 0.54 | 1.87 | 8.38E-03 | 2.71E-02 | Welsh onion | FDR < 0.05, P < 0.05 |
| bin23 | Gammaproteobacteria | Xanthomonadales | Xanthomonadaceae | *Pseudoxanthomonas* _A | - | 0.262 | 0.160 | 0.413 | 0.278 | 0.073 | 0.222 | 0.099 | 0.083 | 0.135 | 0.044 | 0.48 | 2.06 | 1.42E-01 | 1.94E-01 | | |
| bin89 | Alphaproteobacteria | Sphingomonadales | Sphingomonadaceae | *Sphingomonas* | *Sphingomonas mali* | 0.353 | 0.541 | 0.700 | 0.531 | 0.100 | 0.383 | 0.236 | 0.143 | 0.254 | 0.070 | 0.48 | 2.09 | 9.57E-02 | 1.32E-01 | | |
| bin67 | Alphaproteobacteria | Rhizobiales | Rhizobiaceae | *Ensifer* _A | - | 0.438 | 0.672 | 0.787 | 0.632 | 0.103 | 0.276 | 0.287 | 0.330 | 0.298 | 0.017 | 0.47 | 2.12 | 4.36E-02 | 7.16E-02 | Welsh onion | P < 0.05 |
| bin90 | Actinomycetia | Mycobacteriales | Mycobacteriaceae | *Mycobacterium* | *Mycobacterium mucogenicum* | 0.756 | 0.291 | 0.979 | 0.675 | 0.203 | 0.249 | 0.431 | 0.220 | 0.300 | 0.066 | 0.44 | 2.25 | 1.76E-01 | 2.33E-01 | | |
| bin36 | Gammaproteobacteria | Xanthomonadales | Xanthomonadaceae | *Pseudoxanthomonas* _A | - | 0.273 | 0.297 | 0.594 | 0.388 | 0.103 | 0.245 | 0.112 | 0.110 | 0.156 | 0.045 | 0.40 | 2.49 | 6.32E-02 | 9.66E-02 | | |
| bin52 | Alphaproteobacteria | Rhizobiales | Kaistiaceae | *Bauldia* | - | 0.701 | 0.676 | 0.587 | 0.655 | 0.035 | 0.208 | 0.257 | 0.248 | 0.238 | 0.015 | 0.36 | 2.75 | 3.19E-04 | 6.54E-03 | Welsh onion | FDR < 0.05, P < 0.05 |
| bin96 | Gammaproteobacteria | Xanthomonadales | Rhodanobacteraceae | *Dyella* | - | 1.647 | 0.834 | 1.437 | 1.306 | 0.244 | 0.497 | 0.194 | 0.387 | 0.359 | 0.089 | 0.27 | 3.64 | 2.21E-02 | 4.56E-02 | Welsh onion | FDR < 0.05, P < 0.05 |
| bin48 | Alphaproteobacteria | Rhizobiales | Rhizobiaceae | *Rhizobium* _A | *Rhizobium rhizoryzae* | 0.232 | 0.218 | 0.285 | 0.245 | 0.020 | 0.080 | 0.053 | 0.064 | 0.066 | 0.008 | 0.27 | 3.73 | 1.17E-03 | 9.22E-03 | Welsh onion | FDR < 0.05, P < 0.05 |
| bin119 | Alphaproteobacteria | Sphingomonadales | Sphingomonadaceae | *Novosphingobium* | *Novosphingobium aromaticivorans* | 0.262 | 0.435 | 0.473 | 0.390 | 0.065 | 0.123 | 0.090 | 0.095 | 0.103 | 0.010 | 0.26 | 3.79 | 8.63E-03 | 2.72E-02 | Welsh onion | FDR < 0.05, P < 0.05 |
| bin28 | Gammaproteobacteria | Burkholderiales | Burkholderiaceae | *Herbaspirillum* | *Herbaspirillum seropedicae* | 0.330 | 0.238 | 0.293 | 0.287 | 0.027 | 0.062 | 0.097 | 0.059 | 0.073 | 0.012 | 0.25 | 3.95 | 2.79E-03 | 1.44E-02 | Welsh onion | FDR < 0.05, P < 0.05 |
| bin84 | Gammaproteobacteria | Burkholderiales | Burkholderiaceae | *Acidovorax* _F | *Acidovorax monticola* | 0.238 | 0.340 | 0.247 | 0.275 | 0.033 | 0.068 | 0.080 | 0.058 | 0.069 | 0.006 | 0.25 | 4.00 | 9.85E-04 | 9.22E-03 | Welsh onion | FDR < 0.05, P < 0.05 |
| bin38 | Gammaproteobacteria | Burkholderiales | Burkholderiaceae | *Rhizobacter* | - | 0.124 | 0.317 | 0.462 | 0.301 | 0.098 | 0.069 | 0.081 | 0.065 | 0.072 | 0.005 | 0.24 | 4.20 | 7.72E-02 | 1.10E-01 | | |
| bin100 | Gammaproteobacteria | Xanthomonadales | Xanthomonadaceae | *Lysobacter* _B | - | 0.127 | 0.224 | 0.336 | 0.229 | 0.061 | 0.071 | 0.045 | 0.043 | 0.053 | 0.009 | 0.23 | 4.34 | 2.21E-02 | 4.56E-02 | Welsh onion | FDR < 0.05, P < 0.05 |
| bin74 | Unclassified | - | - | - | - | 0.131 | 0.202 | 0.208 | 0.180 | 0.025 | 0.042 | 0.042 | 0.035 | 0.040 | 0.002 | 0.22 | 4.49 | 5.50E-03 | 2.00E-02 | Welsh onion | FDR < 0.05, P < 0.05 |
| bin122 | Unclassified | - | - | - | - | 1.917 | 1.689 | 1.523 | 1.710 | 0.114 | 0.377 | 0.396 | 0.368 | 0.380 | 0.008 | 0.22 | 4.50 | 8.60E-04 | 9.22E-03 | Welsh onion | FDR < 0.05, P < 0.05 |
| bin42 | Gammaproteobacteria | Xanthomonadales | Rhodanobacteraceae | *Dyella* | - | 0.344 | 0.514 | 0.993 | 0.617 | 0.194 | 0.190 | 0.101 | 0.103 | 0.132 | 0.029 | 0.21 | 4.69 | 2.16E-02 | 4.56E-02 | Welsh onion | FDR < 0.05, P < 0.05 |
| bin54 | Alphaproteobacteria | Sphingomonadales | Sphingomonadaceae | *Sphingopyxis* | *Sphingopyxis* sp. 001468395 | 0.393 | 0.447 | 0.573 | 0.471 | 0.053 | 0.110 | 0.091 | 0.079 | 0.093 | 0.009 | 0.20 | 5.05 | 4.48E-04 | 7.87E-03 | Welsh onion | FDR < 0.05, P < 0.05 |
| bin57 | Alphaproteobacteria | Sphingomonadales | Sphingomonadaceae | *Novosphingobium* | - | 0.196 | 0.450 | 0.280 | 0.309 | 0.075 | 0.067 | 0.061 | 0.055 | 0.061 | 0.003 | 0.20 | 5.06 | 2.02E-02 | 4.51E-02 | Welsh onion | FDR < 0.05, P < 0.05 |
| bin123 | Gammaproteobacteria | - | - | - | - | 0.033 | 0.419 | 0.125 | 0.192 | 0.116 | 0.017 | 0.059 | 0.032 | 0.036 | 0.012 | 0.19 | 5.32 | 2.12E-01 | 2.71E-01 | | |
| bin50 | Gammaproteobacteria | Pseudomonadales | Moraxellaceae | *Acinetobacter* | *Acinetobacter pittii* | 0.000 | 0.193 | 0.264 | 0.152 | 0.079 | 0.014 | 0.065 | 0.000 | 0.026 | 0.020 | 0.17 | 5.84 | 4.63E-01 | 5.22E-01 | | |
| bin33 | Alphaproteobacteria | Rhizobiales | Xanthobacteraceae | *Bradyrhizobium* | - | 6.914 | 2.358 | 2.153 | 3.808 | 1.554 | 0.580 | 0.745 | 0.533 | 0.619 | 0.064 | 0.16 | 6.15 | 3.92E-02 | 6.51E-02 | Welsh onion | P < 0.05 |
| bin30 | Unclassified | - | - | - | - | 1.331 | 0.550 | 0.529 | 0.803 | 0.264 | 0.136 | 0.146 | 0.099 | 0.127 | 0.014 | 0.16 | 6.33 | 1.83E-02 | 4.27E-02 | Welsh onion | FDR < 0.05, P < 0.05 |
| bin51 | Gammaproteobacteria | Xanthomonadales | Rhodanobacteraceae | *Luteibacter* | *Luteibacter* sp. 900119235 | 0.470 | 0.670 | 0.366 | 0.502 | 0.089 | 0.081 | 0.083 | 0.064 | 0.076 | 0.006 | 0.15 | 6.60 | 3.54E-03 | 1.44E-02 | Welsh onion | FDR < 0.05, P < 0.05 |
| bin80 | Gammaproteobacteria | Burkholderiales | Burkholderiaceae | *Acidovorax* | *Acidovorax* sp. 000302535 | 0.221 | 0.333 | 0.559 | 0.371 | 0.099 | 0.051 | 0.060 | 0.046 | 0.052 | 0.004 | 0.14 | 7.09 | 1.56E-02 | 4.07E-02 | Welsh onion | FDR < 0.05, P < 0.05 |
| bin86 | Gammaproteobacteria | Burkholderiales | Burkholderiaceae | *Roseateles* | - | 0.229 | 0.371 | 0.375 | 0.325 | 0.048 | 0.043 | 0.046 | 0.040 | 0.043 | 0.002 | 0.13 | 7.53 | 5.47E-03 | 2.00E-02 | Welsh onion | FDR < 0.05, P < 0.05 |
| bin62 | Unclassified | - | - | - | - | 0.279 | 0.329 | 0.360 | 0.323 | 0.024 | 0.046 | 0.037 | 0.030 | 0.037 | 0.005 | 0.12 | 8.61 | 2.28E-04 | 5.62E-03 | Welsh onion | FDR < 0.05, P < 0.05 |
| bin71 | Unclassified | - | - | - | - | 0.109 | 0.522 | 0.436 | 0.355 | 0.126 | 0.045 | 0.045 | 0.030 | 0.040 | 0.005 | 0.11 | 8.96 | 5.11E-02 | 8.12E-02 | | |
| bin121 | Alphaproteobacteria | Sphingomonadales | Sphingomonadaceae | *Sphingobium* | - | 0.440 | 1.038 | 0.936 | 0.805 | 0.185 | 0.103 | 0.087 | 0.065 | 0.085 | 0.011 | 0.11 | 9.47 | 6.42E-03 | 2.26E-02 | Welsh onion | FDR < 0.05, P < 0.05 |
| bin78 | Gammaproteobacteria | Burkholderiales | Burkholderiaceae | *Ralstonia* | *Ralstonia mannitolilytica* | 3.016 | 1.360 | 1.554 | 1.977 | 0.523 | 0.152 | 0.277 | 0.157 | 0.196 | 0.041 | 0.10 | 10.11 | 2.38E-03 | 1.38E-02 | Welsh onion | FDR < 0.05, P < 0.05 |
| bin35 | Unclassified | - | - | - | - | 0.299 | 1.135 | 0.624 | 0.686 | 0.243 | 0.093 | 0.061 | 0.037 | 0.063 | 0.016 | 0.09 | 10.81 | 1.19E-02 | 3.49E-02 | Welsh onion | FDR < 0.05, P < 0.05 |
| bin15 | Unclassified | - | - | - | - | 0.163 | 0.242 | 0.455 | 0.287 | 0.087 | 0.026 | 0.030 | 0.020 | 0.025 | 0.003 | 0.09 | 11.34 | 1.19E-02 | 3.49E-02 | Welsh onion | FDR < 0.05, P < 0.05 |
| bin81 | Chlamydiia | Chlamydiales | Parachlamydiaceae | *Neochlamydia* | - | 0.140 | 0.126 | 0.160 | 0.142 | 0.010 | 0.000 | 0.037 | 0.000 | 0.012 | 0.012 | 0.09 | 11.46 | 6.30E-02 | 9.66E-02 | | |
| bin103 | Unclassified | - | - | - | - | 1.810 | 1.273 | 2.371 | 1.818 | 0.317 | 0.198 | 0.153 | 0.071 | 0.141 | 0.037 | 0.08 | 12.93 | 3.60E-03 | 1.44E-02 | Welsh onion | FDR < 0.05, P < 0.05 |
| bin43 | Unclassified | - | - | - | - | 0.117 | 0.222 | 0.354 | 0.231 | 0.069 | 0.023 | 0.024 | 0.000 | 0.016 | 0.008 | 0.07 | 14.62 | 2.82E-02 | 5.51E-02 | Welsh onion | P < 0.05 |
| bin32 | Gammaproteobacteria | Burkholderiales | Burkholderiaceae | *Ralstonia* | *Ralstonia* sp. 007997035 | 3.585 | 2.404 | 2.600 | 2.863 | 0.365 | 0.216 | 0.220 | 0.106 | 0.181 | 0.037 | 0.06 | 15.85 | 1.74E-03 | 1.19E-02 | Welsh onion | FDR < 0.05, P < 0.05 |
| bin107 | Unclassified | - | - | - | - | 0.084 | 0.490 | 0.367 | 0.314 | 0.120 | 0.034 | 0.025 | 0.000 | 0.019 | 0.010 | 0.06 | 16.12 | 3.52E-02 | 6.02E-02 | Welsh onion | P < 0.05 |
| bin16 | Unclassified | - | - | - | - | 0.475 | 2.381 | 1.281 | 1.379 | 0.553 | 0.096 | 0.062 | 0.060 | 0.072 | 0.012 | 0.05 | 19.04 | 2.11E-02 | 4.56E-02 | Welsh onion | FDR < 0.05, P < 0.05 |
| bin21 | Unclassified | - | - | - | - | 1.329 | 2.448 | 1.565 | 1.781 | 0.341 | 0.092 | 0.065 | 0.051 | 0.069 | 0.012 | 0.04 | 25.64 | 2.18E-04 | 5.62E-03 | Welsh onion | FDR < 0.05, P < 0.05 |
| bin8 | Unclassified | - | - | - | - | 0.424 | 2.235 | 1.175 | 1.278 | 0.525 | 0.069 | 0.036 | 0.033 | 0.046 | 0.012 | 0.04 | 27.81 | 1.28E-02 | 3.50E-02 | Welsh onion | FDR < 0.05, P < 0.05 |
| bin104 | Gammaproteobacteria | Burkholderiales | Burkholderiaceae | *Burkholderia* | *Burkholderia anthina* | 6.162 | 5.159 | 4.293 | 5.205 | 0.540 | 0.166 | 0.153 | 0.100 | 0.139 | 0.020 | 0.03 | 37.37 | 9.86E-05 | 4.63E-03 | Welsh onion | FDR < 0.05, P < 0.05 |
| bin93 | Gammaproteobacteria | LLD-282 | LLD-282 | - | - | 4.925 | 0.000 | 1.244 | 2.056 | 1.479 | 0.000 | 0.016 | 0.000 | 0.005 | 0.005 | 0.00 | 384.59 | 2.21E-01 | 2.80E-01 | | |
| bin120 | Gammaproteobacteria | Pseudomonadales | - | - | - | 0.353 | 0.170 | 0.350 | 0.291 | 0.060 | 0.000 | 0.000 | 0.000 | 0.000 | 0.000 | 0.00 | NA | 3.43E-03 | 1.44E-02 | Welsh onion | FDR < 0.05, P < 0.05 |
| bin65 | Gammaproteobacteria | Pseudomonadales | Spongiibacteraceae | - | - | 0.069 | 0.245 | 0.187 | 0.167 | 0.052 | 0.000 | 0.000 | 0.000 | 0.000 | 0.000 | 0.00 | NA | 1.14E-02 | 3.49E-02 | Welsh onion | FDR < 0.05, P < 0.05 |

The unmapped fraction was excluded from this table and accounted for 41.2 ± 1.2% in Welsh onion samples and 48.6 ± 2.0% in sweet potato samples (mean ± SE, n = 3).
